# Murepavadin is a broad-spectrum outer membrane permeabiliser

**DOI:** 10.64898/2026.06.11.731566

**Authors:** Vedang S. Deshpande, Sarah K.T. Titcombe, Shen Li, Sophia M.A. Riley, Edward J.A. Douglas, Andrew M. Edwards

**Affiliations:** Centre for Bacterial Resistance Biology, Imperial College London, London SW7 2AY, United Kingdom; Department of Infectious Disease, Imperial College London, London SW7 2AY, United Kingdom

## Abstract

Murepavadin is a *Pseudomonas*-specific antibiotic that targets LPS transport protein LptD. However, whilst the mode of action of murepavadin is well defined, the mechanism by which the drug gains access to LptD remains unresolved. Here, we demonstrate a self-directed uptake mechanism for murepavadin, whereby binding to lipid A induces outer membrane disruption, enabling entry of the antibiotic into the periplasm and access to LptD. Murepavadin-LPS interactions were not specific to *P. aeruginosa*, however, and were found to cause OM disruption across a wide range of Gram-negative bacteria, resulting in increased antibiotic susceptibility. We also discovered that murepavadin-mediated OM disruption sensitised *E. coli* to the membrane attack complex of the complement system. In conclusion, murepavadin is a broad-spectrum membrane permeabiliser, which results in increased bacterial susceptibility to antibiotics and host defences.

## Introduction

Antimicrobial resistance (AMR) is a growing global health crisis, responsible for millions of deaths each year. Among the pathogens of greatest concern are Gram-negative bacteria, including *Pseudomonas aeruginosa*, *Acinetobacter baumannii* and members of the Enterobacterales, which exhibit high levels of both intrinsic and acquired resistance to multiple antibiotic classes (1). The limited development of new antibiotics active against Gram-negative organisms has further exacerbated this challenge, creating an urgent need for agents that exploit novel targets and mechanisms of action (2).

A major contributor to the intrinsic resistance of Gram-negative bacteria is the outer membrane (OM) (3). This highly selective barrier consists of selective porins embedded within an asymmetric lipid bilayer, with an outer leaflet of lipopolysaccharide (LPS), a glycolipid that is stabilised by divalent cations, and an inner leaflet of phospholipids (Fig. 1) (4,5). LPS is a complex molecule which is comprised of a densely packed hydrophobic core, known as lipid A, and a highly hydrophilic sugar chain called O-antigen (4,5). This amphipathic structure imposes a major barrier to the permeation of toxic substances such as antibiotics (4,5). Consequently, numerous antibiotics that are active against Gram-positive bacteria fail to accumulate within Gram-negative cells at therapeutically relevant concentrations (4,5). Understanding how antimicrobial compounds overcome this barrier is therefore central to the development of new therapies against multidrug-resistant Gram-negative pathogens (4,5).

**Figure 1.**
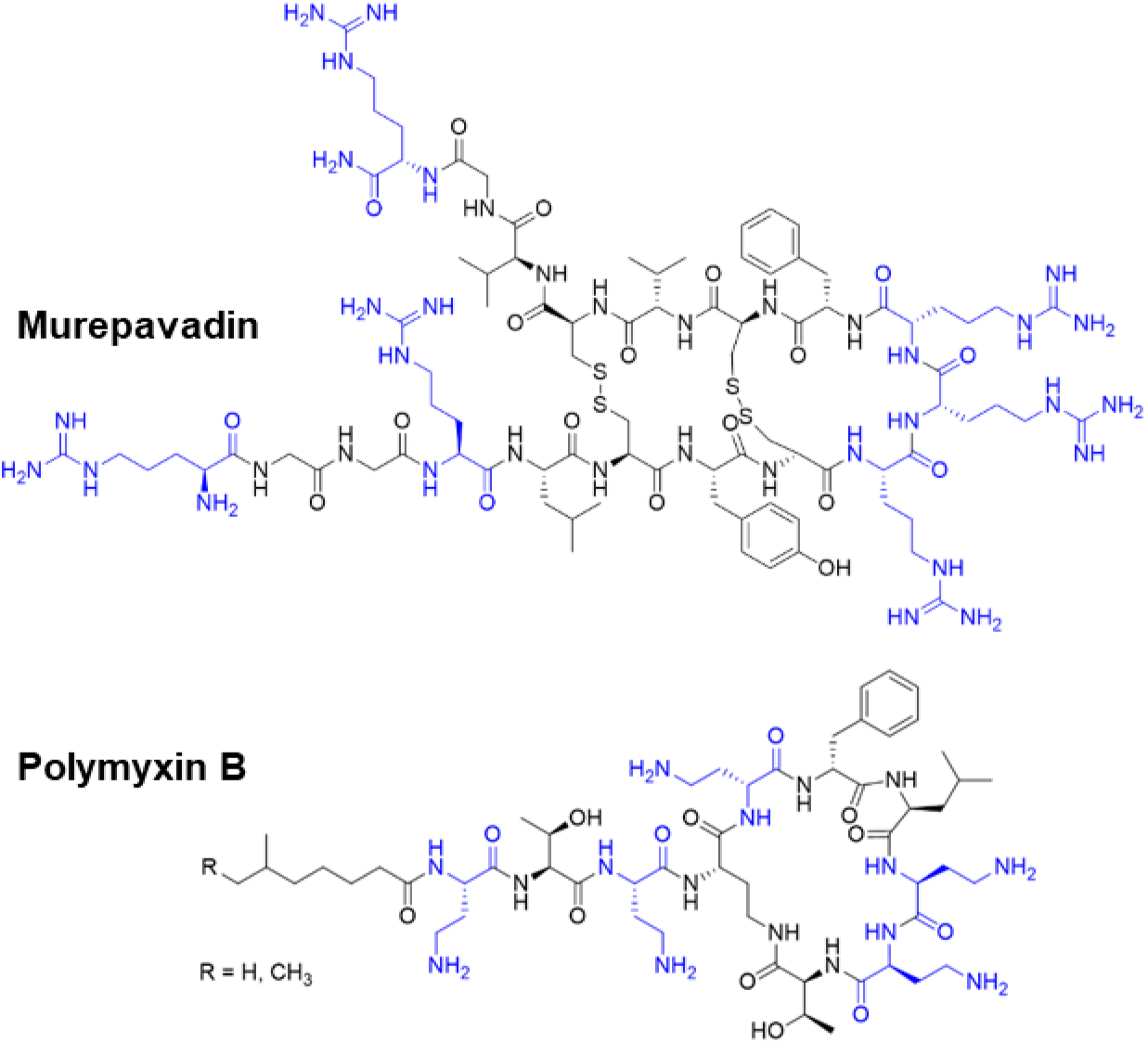
The structures of murepavadin and Polymyxin. **B.** Cationic residues are highlighted in blue. Figure was made using ChemDraw 23.01.

Murepavadin is a synthetic cyclic peptidomimetic antibiotic with potent and highly selective activity against the LPS transport protein LptD of *P. aeruginosa* (6). Analysis of large panels of clinical isolates revealed high levels of susceptibility to murepavadin, including extensively drug-resistant strains (7,8). Murepavadin was subsequently advanced into clinical testing for the treatment of hospital-acquired and ventilator-associated pneumonia via intravenous delivery (9). More recently, inhaled formulations with reduced systemic toxicity are being evaluated for chronic *P. aeruginosa* infections associated with cystic fibrosis and non-cystic fibrosis bronchiectasis (10).

Whilst the target of murepavadin is established, it has not been determined how the antibiotic accesses its binding site on the periplasmic-facing surface of LptD. Structurally, murepavadin is a cyclic β-hairpin peptidomimetic based on the host-defence peptide protegrin-1 (11). It shares notable features with protegrin-1 and the peptide antibiotic polymyxin B (PmB) (11,12). In particular, it possesses multiple positively charged diaminobutyric acid (Dab) residues, making it highly cationic (Fig. 1) (12). For PmB, these Dab residues mediate the electrostatic interaction with the negatively charged phosphate residues of LPS, resulting in displacement of stabilising cations and disruption of OM integrity, enabling the antibiotic to reach the inner membrane, the disruption of which is critical for bacterial killing, the disruption (12–14). The presence of Dab residues in murepavadin led us to hypothesise that this antibiotic accesses its periplasmic target via interactions with LPS, resulting in a loss of OM barrier function that enables antibiotic uptake across the OM.

## Results

### Murepavadin binds LPS from *P. aeruginosa* and *E. coli*

To determine whether murepavadin interacts with LPS, we compared the MICs of murepavadin and PmB against *P. aeruginosa* in the presence or absence of increasing concentrations of purified *P. aeruginosa* LPS. If an antibiotic binds LPS, the addition of exogenous LPS should provide an excess of binding sites, thereby sequestering the antibiotic, reducing its interaction with the bacterial cell surface, and diminishing its antimicrobial activity (15). As expected, and previously reported (16), supplementation of the media with LPS increased the MIC of PmB from 0.125 µg ml^-1^ to >4 µg ml^-1^ in a dose-dependent manner (Fig. 2a). Similarly, the MIC of murepavadin increased from 0.0625 µg ml^-1^ in the absence of LPS to >4 µg ml^-1^ in the presence of 128 µg ml^-1^ LPS. Conversely, the MIC of the non-LPS-binding antibiotic meropenem did not change with LPS supplementation (Supplementary Fig. S1).

**Figure 2.**
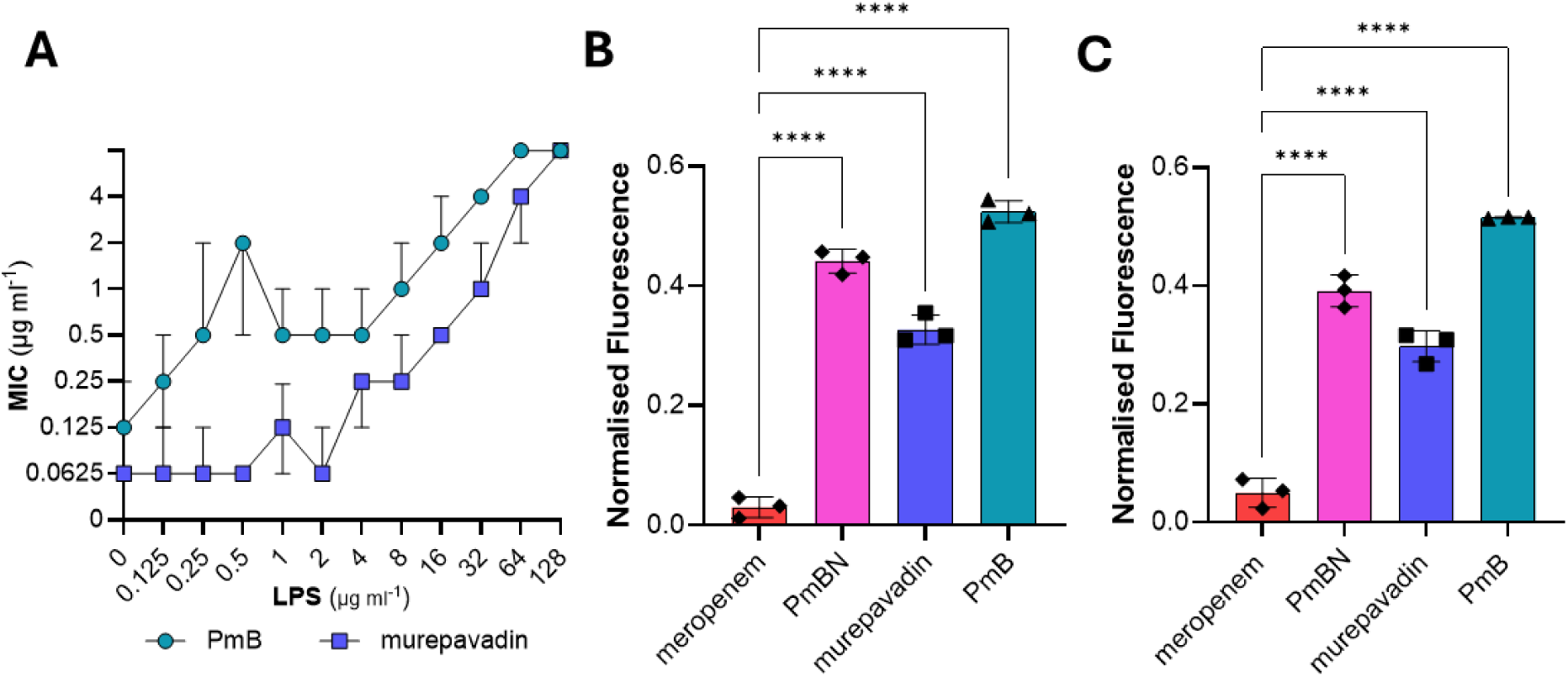
Murepavadin binds LPS. **A)** MIC of PmB and murepavadin against *P. aeruginosa,* in the presence of increasing concentrations of exogenous purified LPS. **B)** Bodipy-cadaverine displacement assay of *P. aeruginosa* exposed, or not, to either 4 µg ml^−1^ meropenem, murepavadin, PmB, or PmBN in MM+G. **C)** Bodipy-cadaverine displacement assay of *E. coli* MG1655 exposed, or not, to either 4 µg ml^−1^ meropenem, murepavadin, PmB, or PmBN in MM+G. Fluorescence emission at _516nm_ was normalised according to OD_600nm_. For B and C, values were blanked according to the untreated control. All experiments were replicated in *n* = 3 independent assays. For A, error bars show the 95% confidence interval of the median. For B and C, error bars show the standard deviation of the mean. For B and C, significant differences were determined between untreated and antibiotic conditions by one-way ANOVA. \**P* < 0.05; \*\**P* < 0.01; \*\*\**P* < 0.001; \*\*\*\**P* < 0.0001.

The Dab residues of PmB mediate an electrostatic interaction with the phosphate moieties of the lipid A component of LPS (12,13,17). As murepavadin also harbours Dab residues, we hypothesised that murepavadin might also bind to the phosphates of lipid A. To assess the interaction between murepavadin and PmB with lipid A, we utilised the fluorescent probe BODIPY-cadaverine. The cadaverine component of this probe specifically binds to the phosphate group of lipid A, resulting in fluorescence quenching of the BODIPY fluorophore. Antibiotics that bind lipid A displace BODIPY-cadaverine, resulting in increased fluorescence. Confirming this, when *P. aeruginosa* was exposed to either PmB or the non-lethal PmB nonapeptide, increased BODIPY-cadaverine fluorescence was observed (Fig. 2b). A similar increase in fluorescence was observed with murepavadin exposure, but not with meropenem (Fig. 2b), confirming that murepavadin binds lipid A.

Lipid A is highly conserved across Gram-negative bacterial species (18). Therefore, we hypothesised that murepavadin may interact with the lipid A of other Gram-negative species. To test this, the BODIPY-cadaverine displacement assay was repeated with *Escherichia coli* MG1655. A similar fluorescence profile was observed, with increased fluorescence upon PmB, PmBN, and murepavadin exposure, but not with meropenem exposure (Fig. 2c). Together, these results show that although murepavadin only displays antimicrobial activity towards *P. aeruginosa,* it can bind to the LPS of other Gram-negative species.

### Murepavadin disrupts the cation bridges of LPS

When PmB binds lipid A, it results in disruption of the cation bridges that stabilise neighbouring LPS molecules. As such, the MIC of PmB is known to increase when excess divalent cations are present (Fig. 3a) (19). Like PmB, supplementation of the media with excess Mg^2+^ increased the MIC of murepavadin against *P. aeruginosa,* indicating that it also disrupts the cation bridges that contribute to OM stability (Fig. 3a).

**Figure 3.**
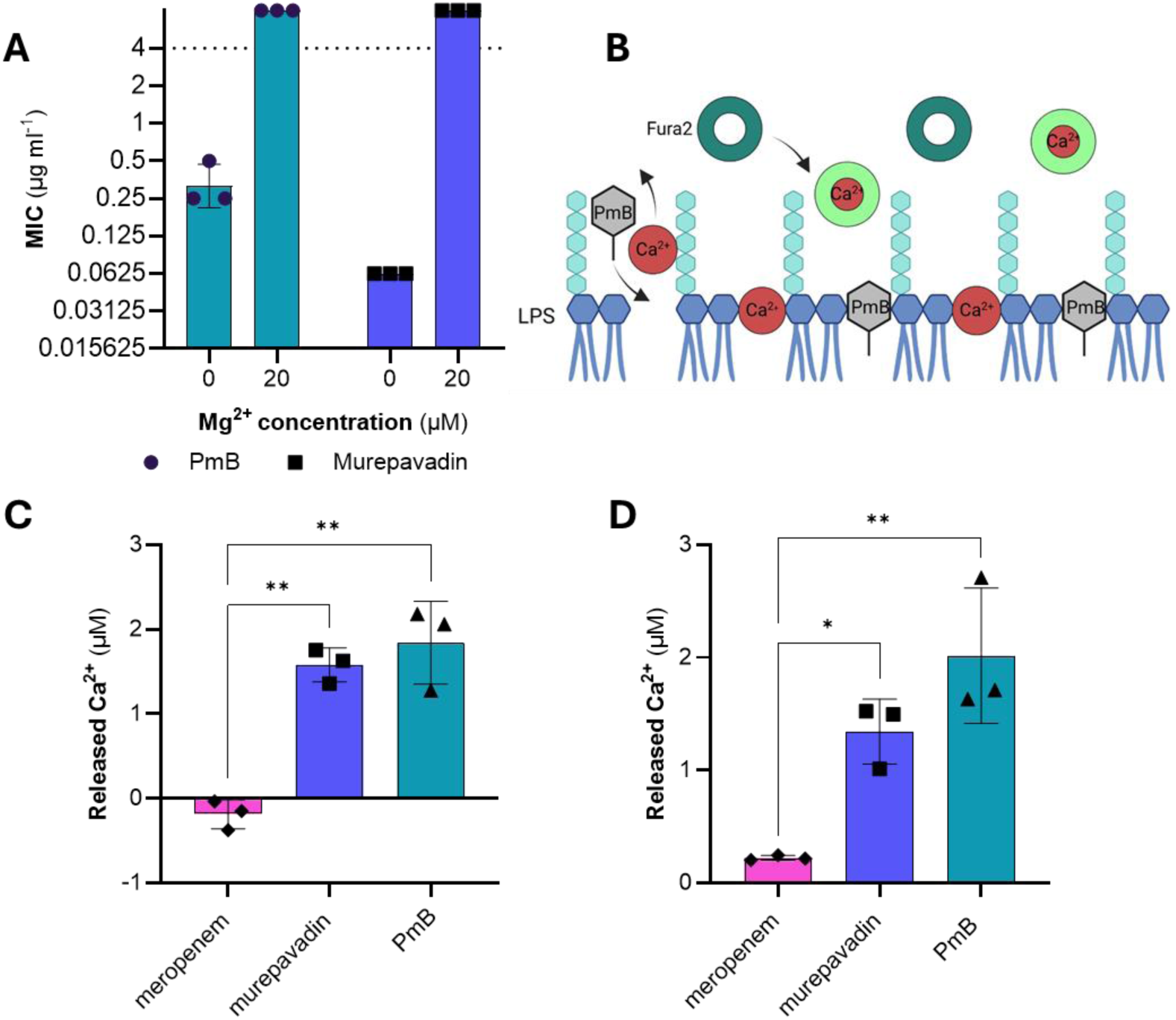
Murepavadin disrupts cationic bridges. **A)** MIC of PmB and murepavadin against *P. aeruginosa,* with and without supplementation of 20 µM Mg^2+^. **B)** Schematic for the detection of cationic bridge disruption using Fura 2. PmB binds to lipid A, resulting in the release of divalent cations into the supernatant. Fura 2 binds to released Ca^2+^, resulting in fluorescence (as indicated by a colour change). **C)** Cation bridge disruption of *P. aeruginosa* cells exposed to 8 µg ml^−1^ meropenem, murepavadin, or PmB, as determined by fluorescence of Fura 2. **D)** Cation bridge disruption of *E. coli* MG1655 cells exposed, or not, to 8 µg ml^−1^ meropenem, murepavadin, or PmB, as determined by fluorescence of Fura 2. For C and D, values were blanked according to the untreated control to give the amount of released Ca^2+^. All experiments were replicated in *n* = 3 independent assays. Error bars show the standard deviation of the mean. For C and D, significant differences were determined between untreated and antibiotic conditions by one-way ANOVA. ‘ns’ not significant; \**P* < 0.05; \*\**P* < 0.01; \*\*\**P* < 0.001; \*\*\*\**P* < 0.0001.

To directly confirm this mechanism, we measured the release of divalent cations following antibiotic treatment by repurposing the ratiometric fluorescent dye Fura-2, a Ca²⁺-binding probe widely used to measure intracellular calcium concentrations (Fig. 3b) (20). A standard curve relating the Fura-2 fluorescence ratio to a serial dilution of CaCl₂ was first generated to enable interpolation of fluorescence measurements (Supplementary Fig. S2) Using this assay, we observed a significant increase in released Ca²⁺ following treatment of *P. aeruginosa* with murepavadin or the LPS-binding control antibiotic PmB (Fig. 3c). Parallel OD_600_ measurements demonstrated that there was no bacterial lysis during the assay, ruling out release of intracellular Ca²⁺ (Supplementary Fig. S3). This assay was repeated with *E. coli,* which also yielded a significant increase in free Ca²⁺ following treatment with murepavadin or PmB (Fig. 3d). Together, these results show that murepavadin disrupts cation bridges of both *P. aeruginosa* and *E. coli* and suggest that murepavadin may compromise OM integrity.

### Murepavadin compromises the OM barrier

To test whether murepavadin-mediated cation bridge disruption was sufficient to compromise OM integrity, we measured the hydrolysis of the chromogenic β-lactam nitrocefin by periplasmic β-lactamase as a readout of OM permeabilisation to small, antibiotic-like molecules (21). Since we had previously established that murepavadin can bind to *E. coli* lipid A (Fig. 3d), we performed the nitrocefin assay in the *E. coli* MG1655 background to rule out any impact from the disruption of LptD, which only occurs with *Pseudomonas*.

Upon exposure of *E. coli* MG1655 to PmB, the rate of nitrocefin hydrolysis increased rapidly and reached saturation within 60 min, confirming the potent OM permeabilising activity of PmB (Fig. 4). Conversely, and in keeping with previous findings (22), PmBN exhibited a lower degree of OM permeabilising activity, with nitrocefin hydrolysis only surpassing the untreated control from 150 min onwards (Fig. 4). Murepavadin likewise induced more modest OM permeabilisation than PmB but displayed faster kinetics than PmBN, resulting in an earlier increase in nitrocefin hydrolysis (Fig. 4). These findings demonstrate that murepavadin-mediated cation bridge disruption is sufficient to compromise OM barrier integrity, sufficient at least to allow the ingress of a small molecule antibiotic analogue.

**Figure 4.**
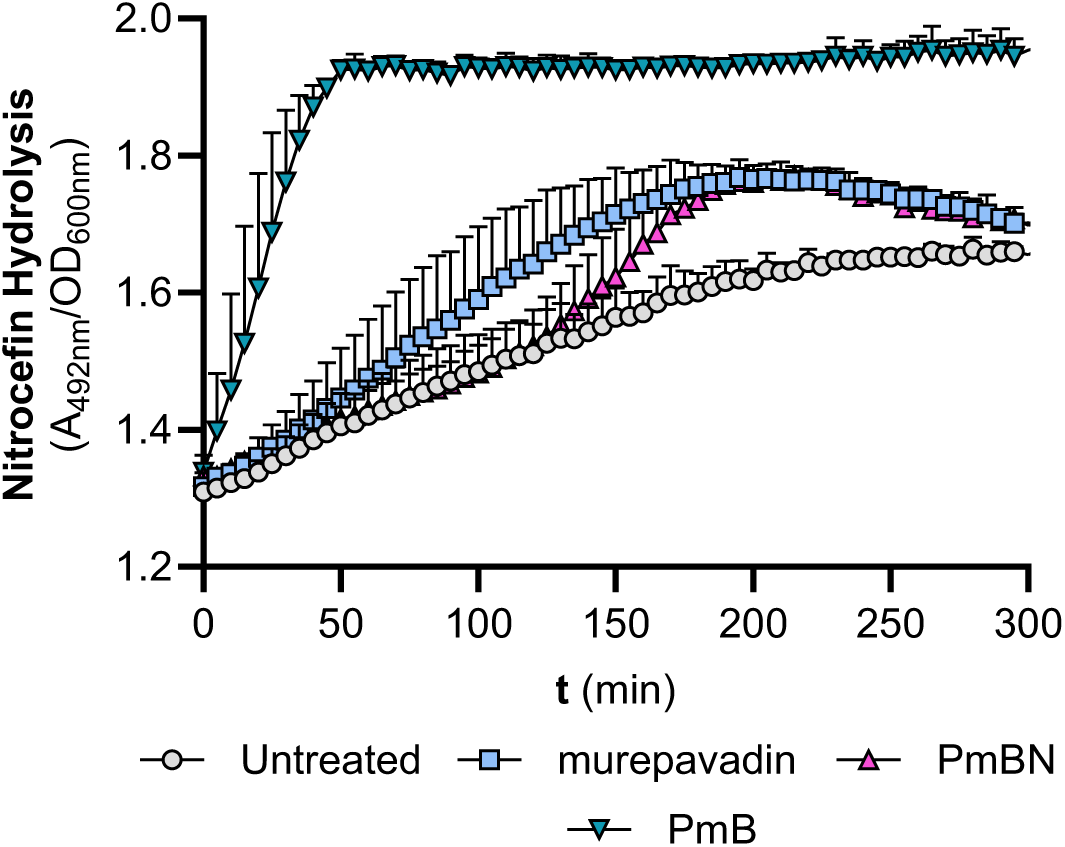
Murepavadin disrupts the OM. OM disruption of *E. coli* MG1655 *mcr1\** cells exposed, or not, to 4 µg ml^−1^ PmB, PmBN, or murepavadin, as determined by hydrolysis or the chromogenic β-lactam nitrocefin. Error bars show the standard deviation of the mean.

### Murepavadin potentiates the growth inhibitory activity of several antibiotics

The OM provides a major barrier to the entry of many antibiotics. Therefore, OM disruption has emerged as a potential strategy for enhancing antibiotic activity (23,24). For example, the non-antimicrobial chelating agent EDTA has been shown to potentiate numerous antibiotics through OM disruption (25). We have also shown recently that EDTA-mediated OM disruption can potentiate PmB activity by enhancing access to LPS in the IM (14).

To test whether murepavadin-mediated OM disruption would enhance the bactericidal activity of PmB, we incubated *E. coli* with murepavadin and a sub-lethal concentration of PmB. The presence of murepavadin enabled PmB-mediated killing via IM disruption, as evidenced by CFU counts and ingress of the SYTOX green nucleic acid stain into the cytoplasm (Fig. 5a, b). Murepavadin did not permeabilise the IM, confirming that the mechanism of bacterial killing was enhanced PmB activity. These experiments were repeated with *P. aeruginosa,* which also showed murepavadin potentiation of PmB activity (Supplementary Fig. S4).

**Figure 5.**
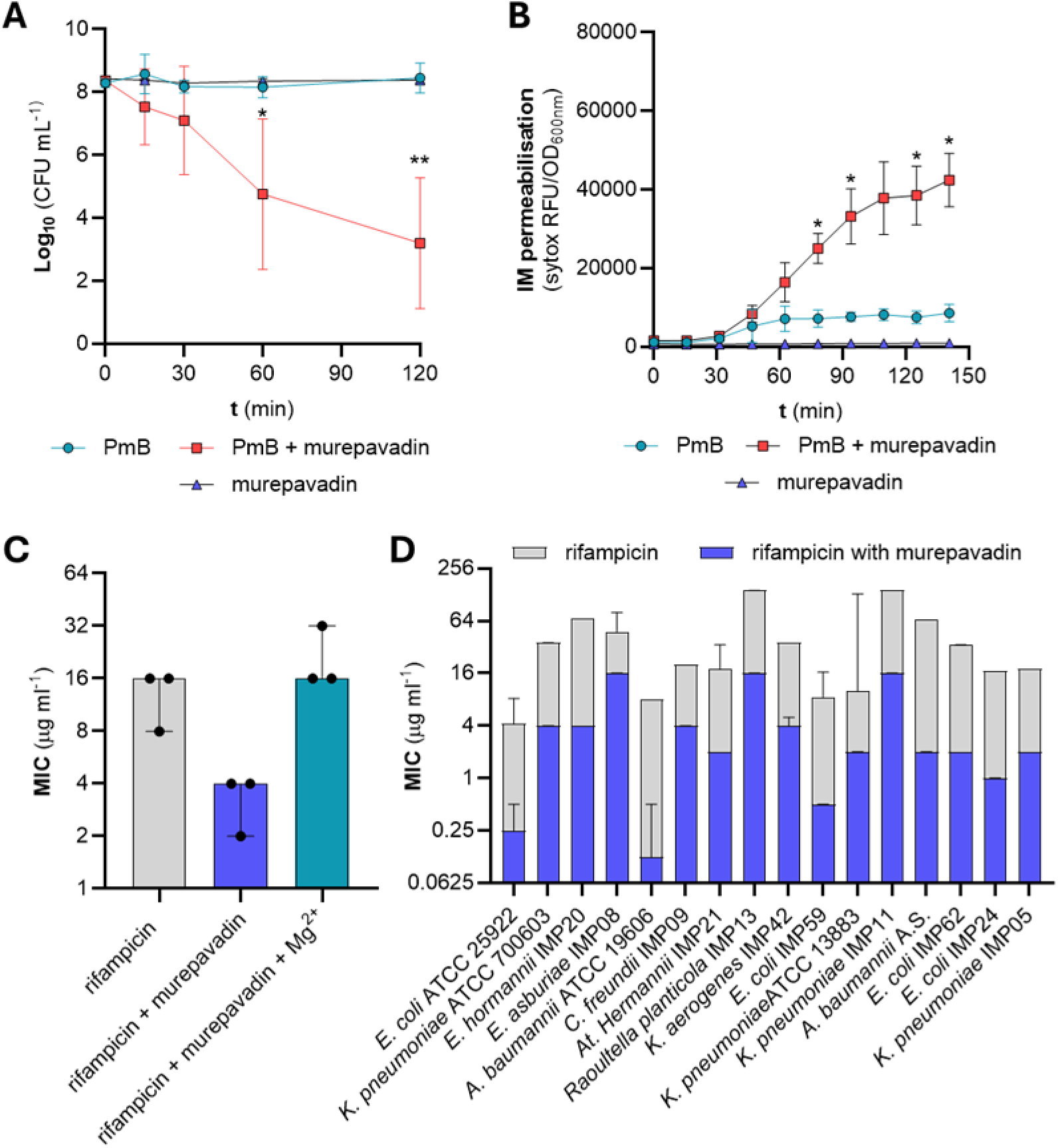
Murepavadin potentiates antibiotics. **A)** Survival of stationary-phase *E. coli* exposed to 4 µg ml^−1^ PmB, with or without 4 µg ml^−1^ murepavadin in MHB, as determined by CFU counts**. B)** Combined OM and IM disruption of stationary-phase *E. coli* exposed to 4 µg ml^−1^ PmB with or without 4 µg ml^−1^ murepavadin in MHB, as determined by uptake of the fluorescent nucleic acid dye SYTOX green. RFU, relative fluorescence units. **C)** MIC of rifampicin with and without murepavadin and 20 µM Mg^2+^, against *E. coli.* **D)** MIC of rifampicin or rifampicin with murepavadin against a range of Gram-negative species. All experiments were replicated in *n* = 3 independent assays. Error bars show the standard deviation of the mean. For A and B, significant differences were determined between PmB and PmB + murepavadin conditions by two-way repeated measures ANOVA. ‘ns’ not significant; \**P* < 0.05; \*\**P* < 0.01; \*\*\**P* < 0.001; \*\*\*\**P* < 0.0001

Having established that murepavadin enhanced PmB penetration into the periplasm, we next investigated whether murepavadin could similarly enhance the activity of large or hydrophobic antibiotics that are otherwise unable to traverse the OM efficiently (Table. 1). In keeping with its OM disrupting activity, murepavadin reduced the MICs of the macrolides azithromycin, clarithromycin, and erythromycin, as well as minocycline, rifampicin, zoliflodacin, PmB, and linezolid, but did not affect vancomycin susceptibility (Table 1).

**Table 1.**
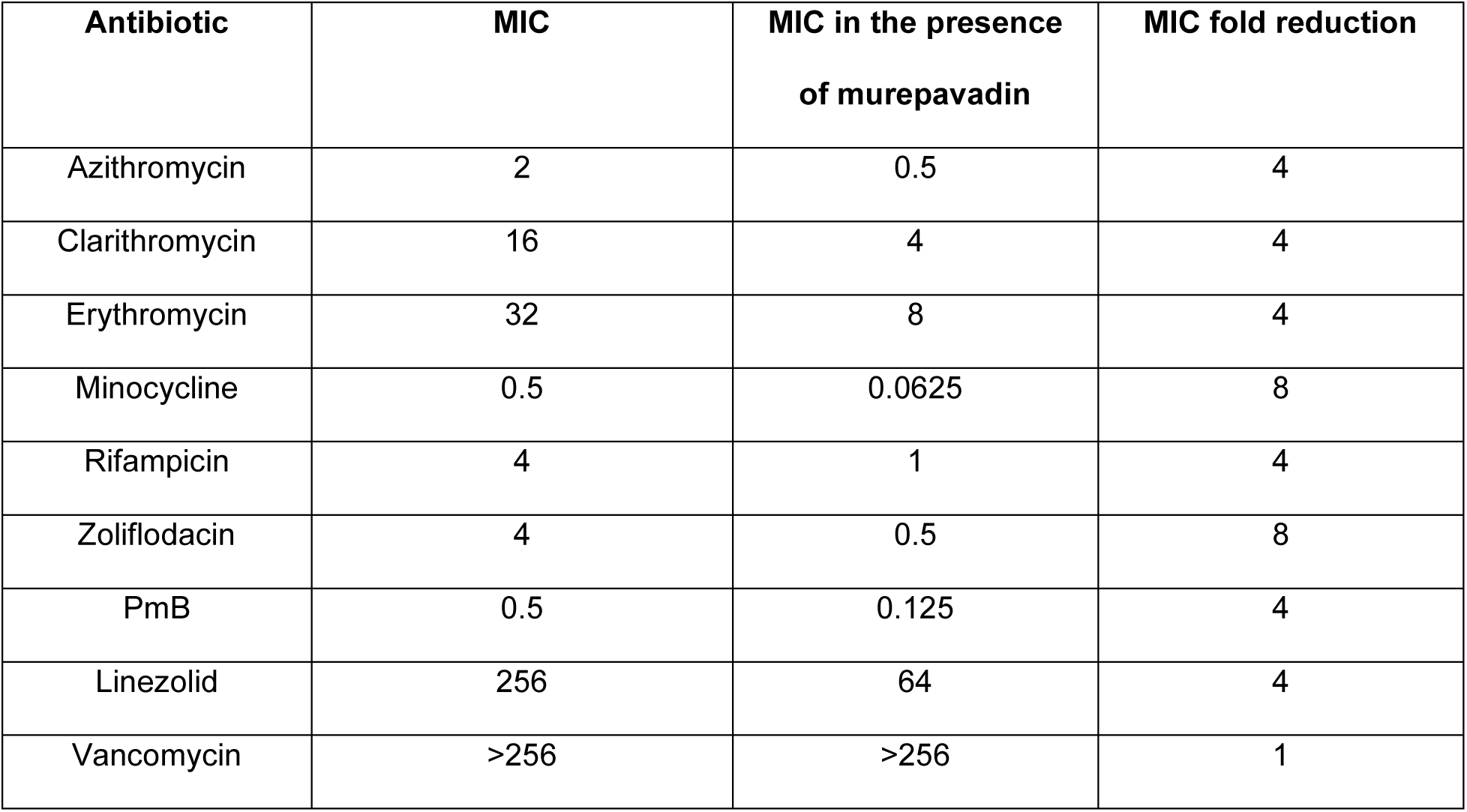
Murepavadin potentiates several antibiotics. Murepavadin was added at a final concentration of 4 µg ml^−1^. All values represent the median of three independent repeats. Note that the highest concentration tested for vancomycin was 256 µg ml^−1^.

PmB-rifampicin is increasingly being used as a salvage therapy for MDR Gram-negative infections (26,27). Likewise, murepavadin potentiated rifampicin activity through a mechanism dependent on cation bridge disruption, as supplementation with excess Mg²⁺ abolished the potentiating effect of murepavadin (Fig. 5c). To examine the clinical potential and spectrum of activity of murepavadin-rifampicin, the MIC of rifampicin was repeated with or without murepavadin against a diverse panel of MDR Gram-negative species (Fig. 5d). In all cases, a reduced rifampicin MIC was observed in the presence of murepavadin, which in most cases was ≥8-fold, with the greatest potentiation observed against *Acinetobacter baumanii* ATCC 19606 (32-fold reduction in rifampicin MIC). Together, these results show that murepavadin is an effective OM permeabiliser which has potential as an antibiotic adjuvant, particularly in combination with rifampicin.

### Murepavadin resensitises *E. coli* to human serum

Previous studies have shown that compounds that permeabilise the OM potentiate bacterial killing by the complement system present in human serum. For example, pre-treatment of serum-resistant Gram-negative bacteria with PmB restores susceptibility to human serum (28). We therefore hypothesised that the OM permeabilising activity of murepavadin may similarly enhance serum-mediated bacterial killing. To test this, we incubated serum-resistant *E. coli* strains CFT073 (Fig. 6a) and CNR1728 (Fig. 6b) with 10% human serum with or without murepavadin. The addition of murepavadin enabled serum-mediated killing against both strains (Fig. 6a, b), as evidenced by a significant reduction in CFU counts. When the killing assay was repeated in heat-inactivated serum, no killing activity was observed, regardless of whether murepavadin was present or not (Supplementary Fig. S5), strongly implying a role for the membrane attack complex (MAC) (29). These findings suggest that the spectrum of activity of murepavadin may extend beyond *Pseudomonas* species when assessed *in vivo*.

**Figure 6.**
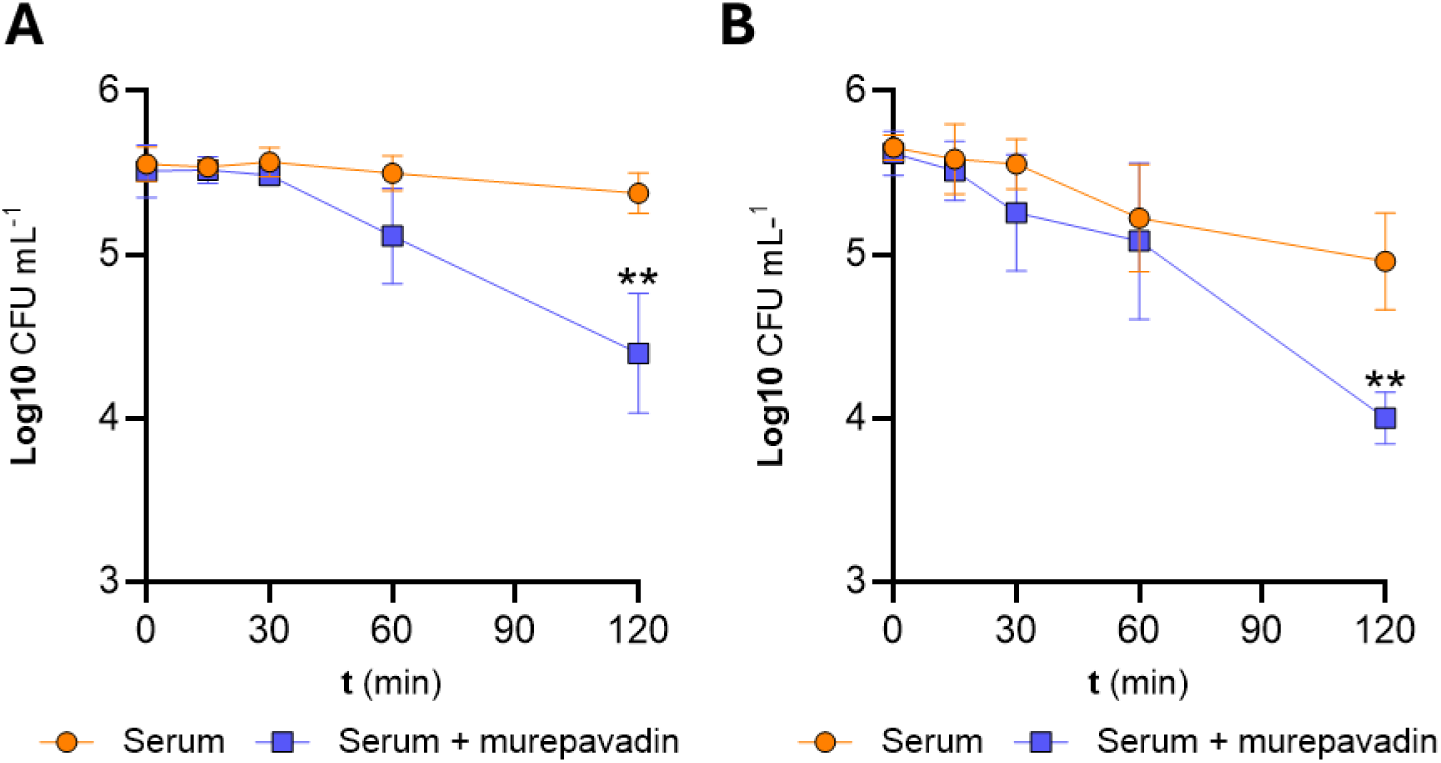
Murepavadin potentiates serum killing. Survival of *E. coli* CFT073 **A)** or *E. coli* CNR1728 **B)** exposed to 10% serum, with or without 4 µg ml^−1^ murepavadin in MHB, as determined by CFU counts. Experiments were replicated in *n* = 3 independent assays. Error bars show the standard deviation of the mean. For A and B, significant differences were determined between conditions by two-way repeated measures ANOVA. \**P* < 0.05; \*\**P* < 0.01

## Discussion

In this study we sought to elucidate the mechanism by which murepavadin reaches its periplasmic target, given that it is too large to traverse OM porins. We hypothesised that murepavadin may exploit a self-promoted uptake mechanism similar to that employed by the peptide antibiotic PmB, owing to their structural similarities (14,30,31). In support of this hypothesis, we have shown that murepavadin binds to LPS, evidenced by decreased susceptibility to the antibiotic in the presence of LPS supplementation and displacement of the lipid A-specific probe BOPIPY-cadaverine. Furthermore, we demonstrate that engagement of LPS by murepavadin disrupts the cation bridges that stabilise neighbouring LPS molecules, as revealed by a novel Fura-2–based assay for detecting this form of OM damage. Importantly, these interactions were not specific to *P. aeruginosa,* as LPS binding and disruption were also observed with *E. coli,* leading us to evaluate murepavadin as a broad-spectrum OM-damaging agent.

While murepavadin has previously been shown to synergise with a range of antibiotics, including PmB, aminoglycosides and β-lactams (32–34), these studies were conducted in *P. aeruginosa*, where the observed synergy was proposed to result from loss of OM barrier function following inhibition of LPS transport to the OM, or the consequent accumulation of LPS in the IM (32–34). In contrast, we demonstrate for the first time that murepavadin treatment enhances the uptake of not only the antibiotic analogue nitrocefin but also a broad range of antibiotics in *E. coli*. As murepavadin does not target the *E. coli* LptD protein, this potentiation cannot be explained by inhibition of LPS transport but instead is consistent with its direct LPS-binding and OM-disrupting activity. We therefore propose a model of self-promoted murepavadin uptake (Fig. 7).

**Figure 7.**
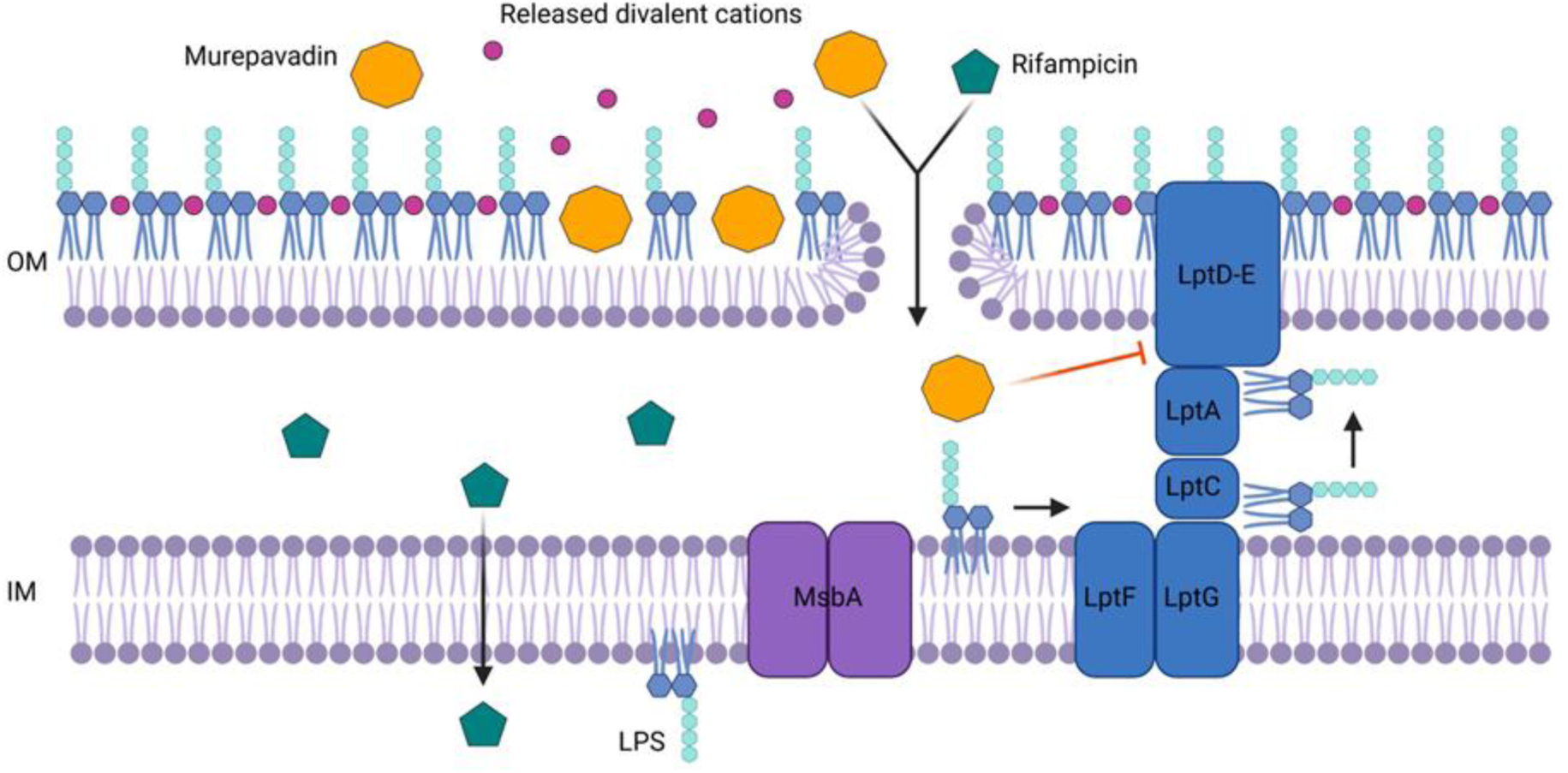
Proposed model for the self-promoted uptake of murepavadin. LPS is synthesised in the cytoplasm before it is flipped into the outer leaflet of the IM by the MsbA protein. LPS is then transported to the outer leaflet of the OM via the Lpt protein complex (blue), which is composed of seven proteins that form a trans-envelope bridge. Murepavadin is electrostatically attracted to LPS through its positively charged Dab residues. Upon binding to lipid A, it disrupts the divalent-cation bridges that stabilise adjacent LPS molecules, thereby releasing membrane-associated cations into the extracellular environment. This destabilisation compromises OM barrier function and facilitates access to the inner leaflet of the OM. From there, murepavadin can traverse the membrane and engage its periplasmic target, LptD. This damage to the OM is also sufficient to allow the ingress of rifampicin.

Among the antibiotics potentiated by murepavadin, murepavadin–rifampicin was identified as particularly promising. Rifampicin is routinely partnered with secondary antibiotics to treat infections by both Gram-positive and Gram-negative bacteria, including in the context of lung infections (26,27,35). Therefore, the treatment of infections associated with cystic fibrosis or non-cystic fibrosis bronchiectasis may be further optimised through a combined dosing regimen of murepavadin and rifampicin. Importantly, this combination would be expected to work against *K. pneumoniae* and *A. baumannii*, which are major causes of lung infections in hospital settings (36). However, further work is required to determine effective doses and the administration of this novel combination.

In addition to potentiating antibiotic activity, we demonstrate that the OM–damaging effects of murepavadin also sensitise clinical *E. coli* isolates to killing by serum complement via the membrane attack complex. The complement system is a ubiquitous innate immune defence component found distributed throughout the body, including in the cystic fibrosis lung environment (37). These findings raise the possibility that murepavadin may exhibit antimicrobial activity against non-*Pseudomonas* species *in vivo,* regardless of a lack of direct antibacterial activity.

This premise provides additional support for the use of host-mimicking conditions in antibiotic testing and development. Numerous studies have shown that antimicrobial potency diverges from rich laboratory media when tested in *ex vivo* conditions, such as serum, or in media designed to artificially recapitulate host-specific niches (38–42). Our results suggest that this limitation may extend even further, with conventional *in vitro* assays potentially overlooking compounds whose antimicrobial activity is contingent upon interactions with the host and therefore only manifests *in vivo*.

Crucially, this study demonstrates that investigating antibiotic interactions with the OM can provide important insights into their mechanisms of action and uptake. Future studies should prioritise determining whether the phenomena described here extend to other antibiotics that target OM biogenesis. For example, Roche’s late-stage antibiotic candidate zosurabalpin, an *Acinetobacter*-specific inhibitor of the LptB₂FGC transporter, is also cationic, albeit to a lesser extent than PmB or murepavadin (43). Nevertheless, establishing how zosurabalpin traverses the OM may reveal opportunities for extending its spectrum of use.

## Methods

### Bacterial strains and growth conditions

The bacterial strains used in this study are listed in Supplementary Table 1. *E. coli* strains expressing *mcr*-1 or *mcr*-1* under the control of the arabinose-inducible *araBAD* (pBAD) promoter were based on those described previously and were transformed with plasmids synthesised by Thermo Fisher and selected using ampicillin 100 µg ml^-1^. All strains were grown in MHB (Millipore) for 18 h at 37 °C with shaking (180 r.p.m.) to stationary phase. Unless stated otherwise, bacteria were washed three times with 1× M9 minimal medium (Gibco) supplemented with 2 mM MgSO_4_ and 0.1 mM CaCl_2_ (referred to here as minimal medium, MM). Washed cells were then resuspended in MM ± 0.36% glucose (MM + G) to 10^8^ CFU ml^−1^. For the Fura-2 assay, 1x M9 minimal medium was used, but without supplementation of cations. To enumerate bacterial CFU, 10-fold serial dilutions were made in sterile PBS and plated onto Mueller–Hinton agar (MHB supplemented with 1.5% bacteriological agar). Inoculated agar plates were incubated statically for 18 h in air at 37 °C.

**Table 1.** Strains used in this study.

| Strain | Description | Reference |
| --- | --- | --- |
| *P. aeruginosa* PA14 | Well-characterised laboratory strain | (44) |
| *E. coli* MG1655 | Well-characterised K-12 laboratory strain | (45) |
|  | *Pmcr-1\**; MG1655 expressing a catalytically inactive variant of Mcr1. AmpR | (14) |
| *E. coli* ATCC 25922 | Type strain used as a reference for antibiotic susceptibility testing | (46) |
| *E. coli* CNR1728 | Clinical isolate | (47) |
| *E. coli* IMP59 | Clinical isolate | (48) |
| *E. coli* IMP62 | Clinical isolate | (48) |
| *E. coli* IMP24 | Clinical isolate | (48) |
| *E. asburiae* IMP08 | Clinical isolate | (48) |
| *E. hormanii* IMP20 | Clinical isolate | (48) |
| *C. freundii* IMP09 | Clinical isolate | (48) |
| *K. pneumoniae* ATCC 700603 | Clinical isolate | (48) |
| *K. pneumoniae* ATCC 13883 | Clinical isolate | (48) |
| *K. pneumoniae* IMP11 | Clinical isolate | (48) |
| *K. pneumoniae* IMP05 | Clinical isolate | (48) |
| *K. aerogenes* IMP42 | Clinical isolate | (48) |
| *A. baumannii* ATCC 19606 | Clinical isolate | (48) |
| *A. baumannii* A.S. | Clinical isolate | (48) |
| *At. hermannii* IMP21 | Clinical isolate | (48) |
| *R. planticola* IMP13 | Clinical isolate | (48) |

### Determination of MICs

The MICs of PmB, murepavadin, meropenem, and rifampicin against the indicated bacterial strains were determined according to the well-established broth microdilution method (49). Briefly, a 96-well microtiter plate (Corning) was used to perform a range of antibiotic concentrations in MHB via a 2-fold serial dilution. Following this, stationary-phase bacteria were diluted to 1 x 10^6^ CFU ml^-1^, before using 100 µl to seed the microtiter plate and give a final inoculum density of 5 x 10^5^ CFU ml^-1^. Where indicated, purified *P. aeruginosa* LPS at a range of concentrations or 20 µM MgCl_2_ was added to the MHB. Plates were then incubated statically at 37 °C for 18 h in air. At that point, the MIC was defined as the lowest antibiotic concentration at which there was no visible growth of bacteria.

### BODIPY-cadaverine displacement assay

The conjugated BODIPY-cadaverine (Invitrogen) fluorophore was used to quantify the interaction of antibiotics with lipid A (50). Stationary-phase *P. aeruginosa* and *E. coli* were washed five times in MM (Gibco) and concentrated to 2 x 10^8^ CFU ml^-1^ in MM+G, before using 100 µl to seed a black microtitre plate with clear-bottomed wells (Greiner Bio-One), to give a final inoculum density of 1 × 10^8^ CFU ml^−1^. BODIPY-cadaverine was added at a final concentration of 5 μg ml^-1^ in MM+G, giving a total volume of 150 µl. Baseline fluorescence was measured in a Tecan Spark plate reader at an excitation wavelength of 580 nm and an emission wavelength of _616nm_. The plate was taken out and left untreated or treated with 4 μg ml^-1^ PmB, murepavadin, PmBN, or meropenem, yielding a final volume of 200 μl per well. BODIPY-cadaverine fluorescence was quantified using the same wavelengths and apparatus as previously described and normalised according to their OD_600nm_ values.

### Quantification of cation bridge disruption

The ability of antibiotics to disrupt the cation bridges of lipid A was determined by repurposing the ratiometric dye Fura-2 (Invitrogen) (20). Stationary-phase *P. aeruginosa* or *E. coli* was washed five times in 1x M9 without cations and concentrated to 1 x 10^9^ CFU ml^-1^. Following this, 300 µl was added to 3 ml of 1x M9 without cations and supplemented with 0.36% glucose and 8 μg ml^-1^ of PmB, murepavadin, PmBN or meropenem and incubated at 37°C with shaking (180 r.p.m.) for 15 min. Immediately following antibiotic incubation, 200 μl was removed and OD_600nm_ recorded to ensure bacterial lysis had not occurred. Samples were then pelleted at 16000 x *g*, and 100 μl of supernatant was transferred to a black microtitre plate with clear-bottomed wells (Greiner Bio-One). Fura-2 at a final concentration of 0.5 μg ml-1 was added to the plate to give a total volume of 200 μl. Fura-2 fluorescence was measured at excitation wavelengths of 340 nm (Ca^2+^ bound) and 380 nm (Ca^2+^ unbound), and an emission wavelength of 510 nm in the Tecan Spark plate reader. Free Ca^2+^ was calculated using the following formula:

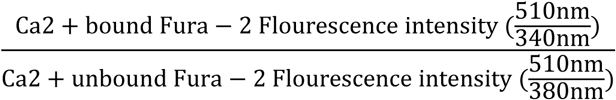

### OM disruption by nitrocefin hydrolysis

To detect OM damage by antibiotics, ingress and hydrolysis of the chromogenic β-lactam nitrocefin were used. Following three washes in MM, stationary-phase *E. coli mcr1\** was diluted to 2 × 10^8^ CFU ml^−1^ in MM+G, and 100 μl was used to seed a 96-well microtiter plate (Corning), to give a final inoculum density of 1 × 10^8^ CFU ml^−1^. Antibiotics were added to each well in a 50 μl volume and at final concentrations of 4 μg ml^-1^ PmB and PmBN, and a range of concentrations of murepavadin; 4-32 μg ml^-1^. Similarly, 50 μl of nitrocefin at a final concentration of 120 μg ml-1 was added to each well, to give a total volume of 200 μl. Nitrocefin hydrolysis was measured every 5 min for 300 min at an absorbance wavelength of _492nm_ in the Tecan Spark plate reader at 37 °C with shaking. Raw absorbance readings were normalised to their OD_600nm_ values.

### Antibiotic time kill assays

Stationary-phase *P. aeruginosa* and *E. coli* were added at a final concentration of 1 × 10^8^ CFU ml^−1^ to 3 ml of MHB and supplemented with PmB (4 µg ml^−1^) or murepavadin (4 µg ml^−1^) when required. For serum time kill assays, stationary-phase *E. coli* CFT073 and CNR1728 were added at a final concentration of 5 × 10^5^ CFU ml^−1^ to 3 ml MHB containing 10% human serum (male type AB, H4522 Sigma Aldrich), with murepavadin (4 µg ml^−1^) when required. Cultures were incubated at 37 °C with shaking (180 r.p.m.). At each time point (0, 15, 30, 60 and 120 min), 20 μl aliquots were taken, serially diluted 10-fold in PBS and plated on MHBA to determine bacterial viability by determination of CFU counts.

### IM permeability assay

To measure IM permeabilisation by PmB, the well-established SYTOX green assay was used (51). Following three washes in MM, stationary-phase *P. aeruginosa* and *E. coli* at inoculum densities of 1 × 10^8^ CFU ml^−1^ were added to 3 ml of MM + G containing PmB (4 µg ml^−1^), murepavadin (4 µg ml^−1^), or both. The fluorescent probe SYTOX green (Invitrogen) was added to these cultures at a final concentration of 1 µM. Aliquots (200 μl) were transferred to a black microtitre plate with clear-bottomed wells (Greiner Bio-One). Fluorescence and OD_600nm_ were measured in a Tecan Spark plate reader (excitation at 535 nm, emission at 617 nm) every 15 min for 2 h at 37 °C with shaking. Raw fluorescence readings were normalised according to OD_600nm_ values.

### Statistics

Experiments were performed on at least three independent occasions, and the resulting data are presented as the arithmetic mean of these biological repeats unless stated otherwise. Error bars, where shown, represent the standard deviation of the mean unless otherwise stated. For multiple comparisons at a single time point or concentration, data were analysed using one-way analysis of variance (ANOVA). When data were obtained at multiple time points or concentrations, two-way ANOVA was used for statistical analysis. Appropriate post hoc tests (Dunnett’s, Tukey’s, Sidak’s, Dunn’s) were carried out to correct for multiple comparisons, with details provided in the figure legends. Asterisks on graphs indicate significant differences between data. All statistical analyses were performed using GraphPad Prism 7 software (GraphPad Software).

## Contributions

V.S.D., S.K.T.T., E.J.A.D., A.M.E. designed the research. V.S.D., S.K.T.T., S.L., S.M.A.R., E.J.A.D., A.M.E. performed the research. V.S.D., S.K.T.T., S.L., S.M.A.R., E.J.A.D., A.M.E. analysed data. V.S.D., S.K.T.T., S.L., S.M.A.R., E.J.A.D., A.M.E. wrote the paper. E.J.A.D., A.M.E. conceptualised the project. E.J.A.D., A.M.E. administered the project. A.M.E. acquired funding. E.J.A.D., A.M.E. contributed equally.

## Corresponding authors

Correspondence to Edward J.A. Douglas, Andrew M. Edwards.

