## Supplementary figures S1-S5 for "Murepavadin is a broad-spectrum outer membrane permeabiliser"

Supplementary data file.

Supplementary figures S1 – S5.

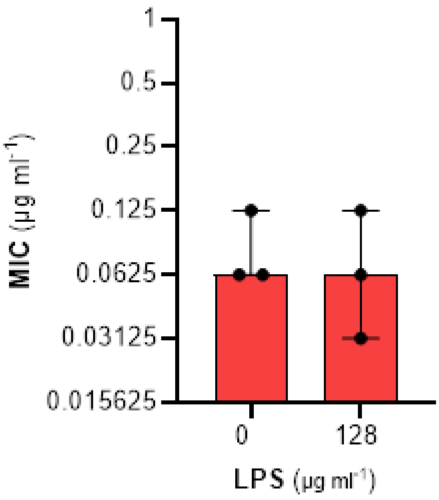


**Supplementary Figure S1. Meropenem does not bind LPS. A)** MIC of meropenem against *P. aeruginosa,* with or without supplementation with exogenous purified LPS. All experiments were replicated in $n$ = 3 independent assays. Error bars show the 95% confidence interval of the median.

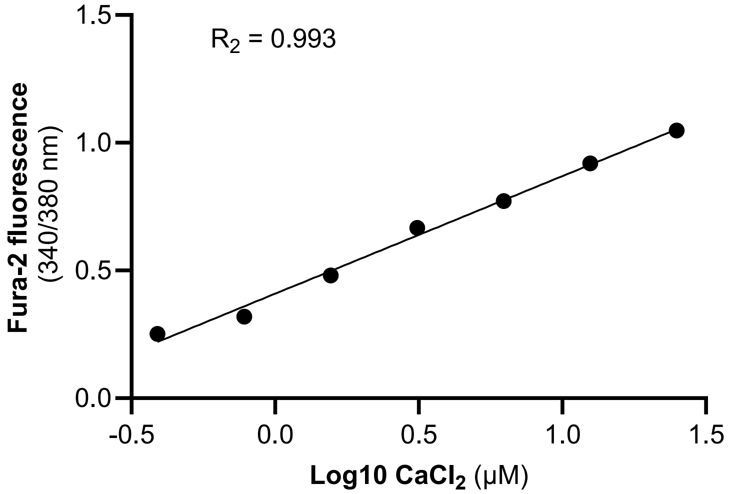


**Supplementary Figure S2. Fura-2 standard curve.** Standard curve plotting $Log_{10}$ $CaCl_2$ concentration (µM) against Fura-2 fluorescence ratio. Simple linear regression was performed using Prism version 10.4.1, with the line representing the best fit. The $R_2$ is shown on the graph.

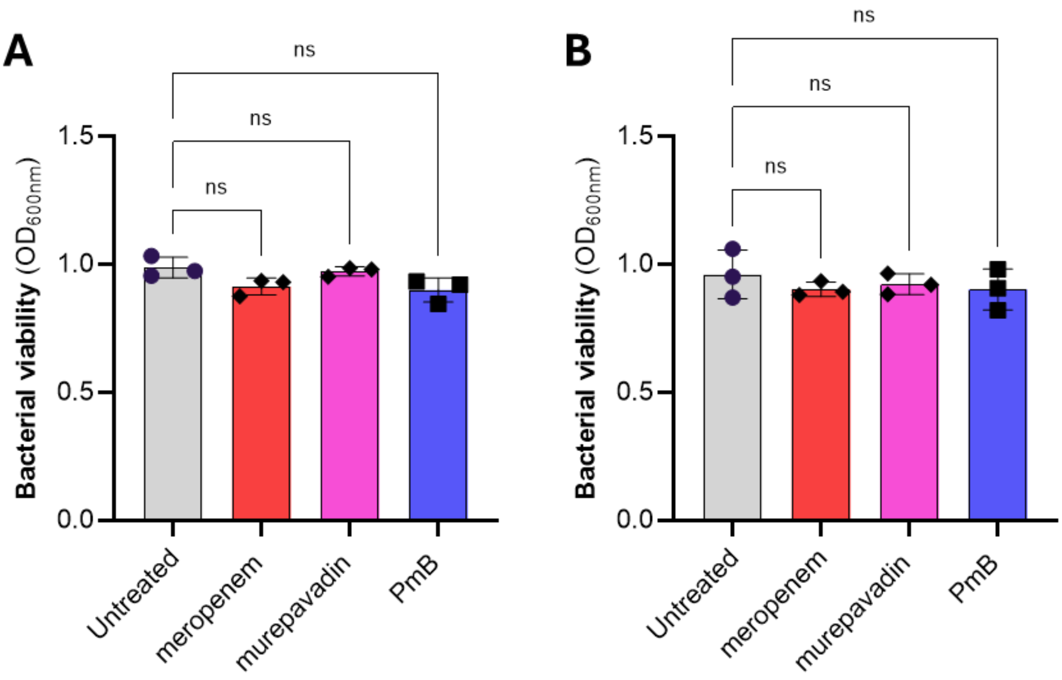


**Supplementary Figure S3. Bacterial lysis is not responsible for increased $Ca^{2+}$ concentration in supernatants. A)** Bacterial viability of *P. aeruginosa* following a 15-minute treatment with 8 µg ml−1 meropenem, murepavadin, or PmB, as determined by $OD_{600nm}$. **B)** Bacterial viability of *E. coli* following a 15-minute treatment with 8 µg ml−1 meropenem, murepavadin, or PmB, as determined by $OD_{600nm}$. All experiments were replicated in *n* = 3 independent assays. Error bars show the standard deviation of the mean. Significant differences were determined between untreated and antibiotic conditions by one-way ANOVA. 'ns' not significant; *$P < 0.05$; **$P < 0.01$; ***$P < 0.001$; ****$P < 0.0001$

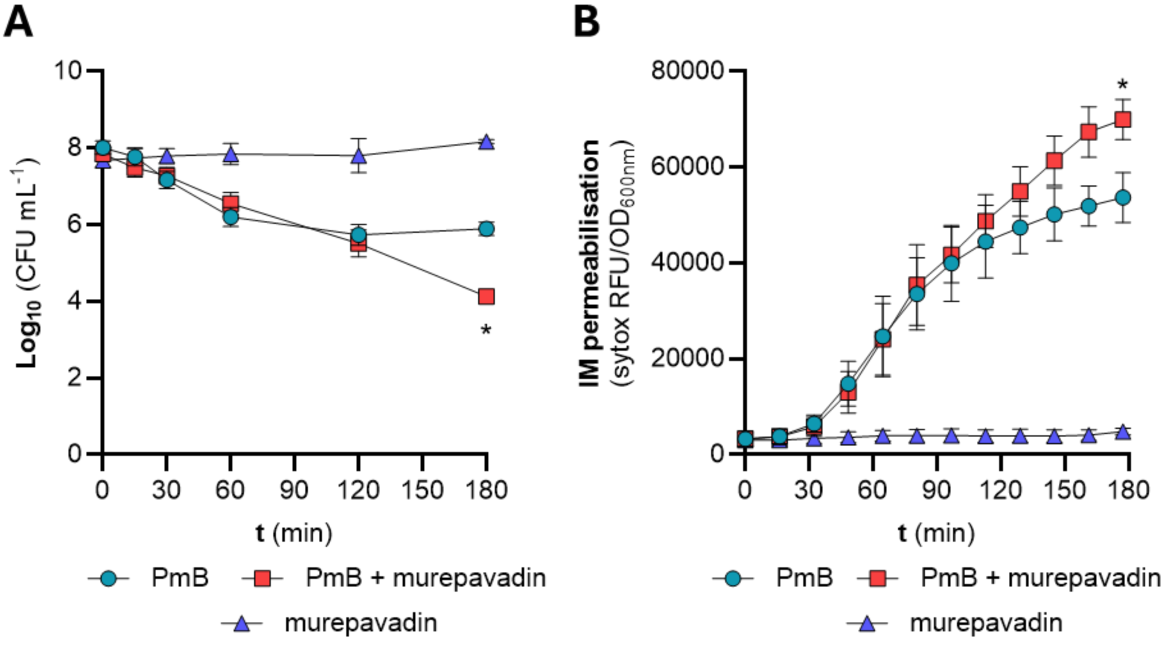


**Supplementary Figure S4. Murepavadin potentiates PmB against *P. aeruginosa*. A)** Survival of stationary-phase *P. aeruginosa* exposed to 2 μg ml$^{-1}$ PmB, with or without 4 μg ml$^{-1}$ murepavadin in MHB, as determined by c.f.u. counts**. B)** Combined OM and IM disruption of stationary-phase *P. aeruginosa* exposed to 2 μg ml$^{-1}$ PmB with or without 4 μg ml$^{-1}$ murepavadin in MHB, as determined by uptake of the fluorescent nucleic acid dye SYTOX green. RFU, relative fluorescence units. All experiments were replicated in $n$ = 3 independent assays. Error bars show the standard deviation of the mean. For A and B, significant differences were determined between PmB and PmB + murepavadin conditions by two-way repeated measures ANOVA. 'ns' not significant; *$P$ < 0.05; **$P$ < 0.01; ***$P$ < 0.001; ****$P$ < 0.0001

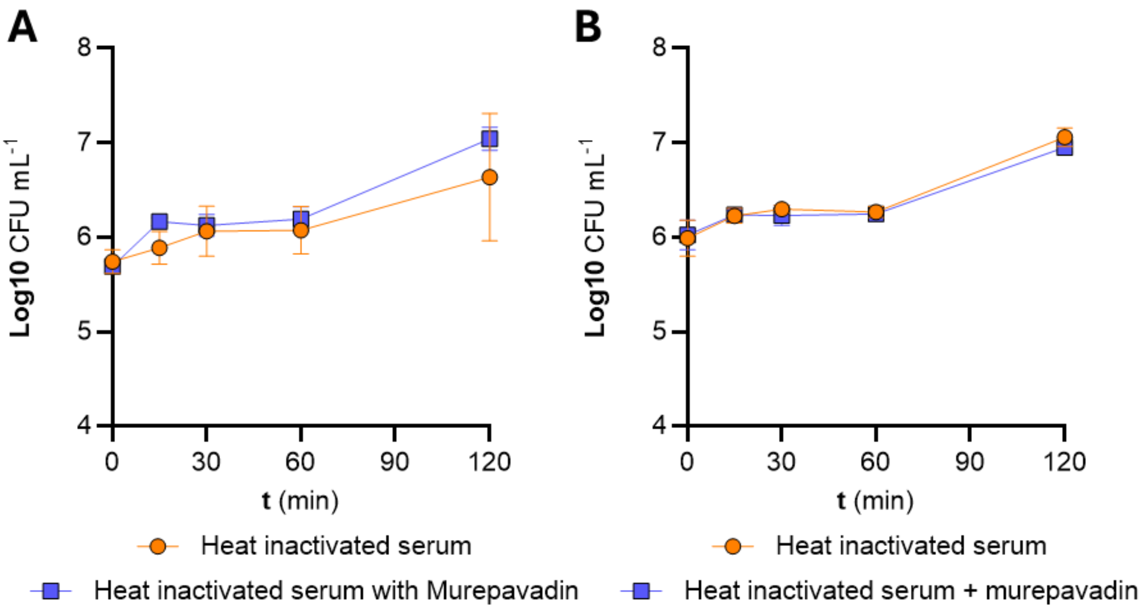


**Supplementary Figure S5. Murepavadin does not promote killing in heat-inactivated serum.** Survival of *E. coli* CFT073 **A)** or *E. coli* CNR1728 **B)** exposed to 10% heat-inactivated serum, with or without 4 µg $ml^{-1}$ murepavadin in MHB, as determined by CFU counts. Experiments were replicated in $n = 3$ independent assays. Error bars show the standard deviation of the mean.
